# Quantitation of FGFR3 signaling via GRB2 recruitment on micropatterned surfaces

**DOI:** 10.1101/2022.04.11.487861

**Authors:** Ingrid Hartl, Veronika Brumovska, Yasmin Striedner, Atena Yasari, Gerhard J. Schütz, Eva Sevcsik, Irene Tiemann-Boege

## Abstract

Fibroblast growth factor receptors (FGFRs) initiate signal transduction via the RAS/MAPK pathway by their tyrosine-kinase activation known to determine cell-growth, tissue differentiation and apoptosis. Recently, many missense mutations have been reported for *FGFR3*, but we only know the functional effect for a handful of them. Some of these mutations result in aberrant FGFR3 signaling and are associated with various genetic disorders and oncogenic conditions. Here we employed micropatterned surfaces to specifically enrich fluorophore-tagged FGFR3 (mGFP-FGFR3) in certain areas of the plasma membrane of living cells. Receptor activation was then quantified via the recruitment of the downstream signal transducer GRB2 tagged with mScarlet (GRB2-mScarlet) to FGFR3 patterns. With this system, we tested the activation of FGFR3 upon ligand addition (fgf1 and fgf2) in the wildtype (WT), as well as in different FGFR3 mutants associated with congenital disorders (G380R, Y373C, K650Q, K650E). Our data showed that the addition of ligands increased GRB2 recruitment to WT FGFR3, with fgf1 having a stronger effect than fgf2. For all mutants, we found an increased basal receptor activity, and only for two of the four mutants (G380R and K650Q), activity was further increased upon ligand addition. Compared to previous reports, two mutant receptors (K650Q and K650E) had either an unexpectedly high or low activation state, respectively. This may be explained by the different receptor populations probed, since the micropatterning method specifically reports on signaling events at the plasma membrane.

**Graphical Abstract:** *Specifications: The maximum size of the image should be 200 × 500 pixels with a minimum resolution of 300 dpi, using Arial font with a size of 10-16 points; Preferred file types: TIFF, EPS, PDF or MS Office files*

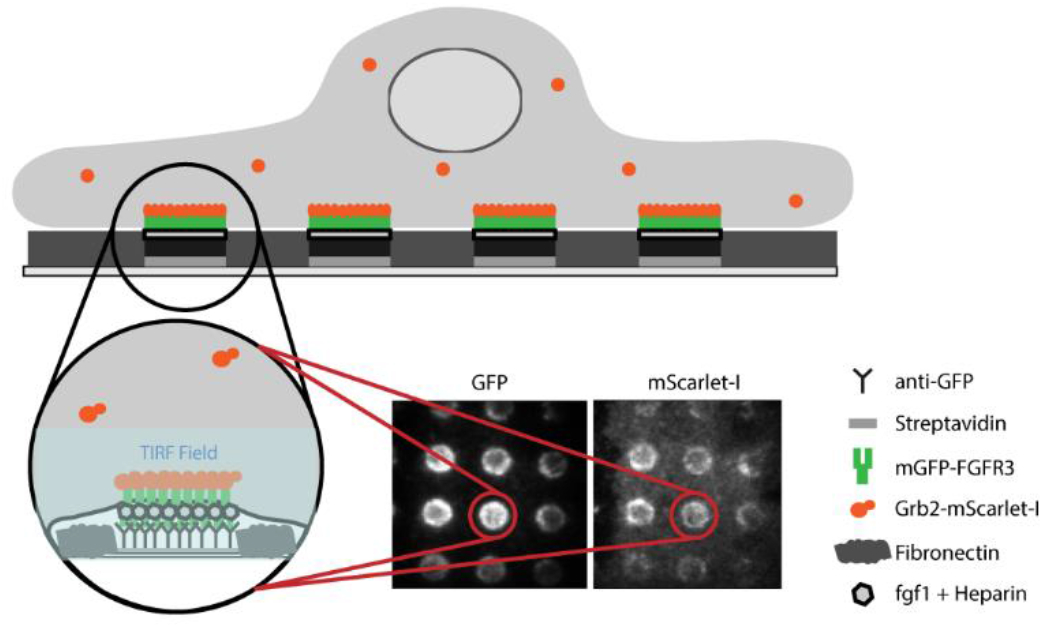

**Research Highlights:**

- Quantification of FGFR3 signaling in live cells on micropatterned surfaces
- Analysis of GRB2 recruitment to the mature receptor at the plasma membrane
- Ligand-independent kinase activation of FGFR3 mutants
- Activation of FGFR3 at the cell surface can be different than in bulk cell extracts

## Introduction

As a family of cell surface receptors, receptor tyrosine kinases (RTKs) initiate intracellular signaling cascades that activate or inhibit different transcription factors linked to numerous cellular processes such as cell growth, development and apoptosis (Lemmon and Schlessinger 2010). Although different RTKs activate the same intracellular signaling pathways, the cellular response can be diverse. Fibroblast growth factor receptors (FGFRs) are a subclass of RTKs and comprise 4 members (Farrell and Breeze 2018) that are evolutionary highly conserved across multi-cellular species (Rebscher et al. 2009; Lemmon and Schlessinger 2010). FGFRs are activated by binding of their ligand fgf, which typically involves receptor dimerization (Ornitz and Itoh 2015), although formation of larger clusters has been observed for FGFR1 in a ligand-specific manner suggesting that the FGFR oligomerization state mediates cellular responses to different ligands (Karl et al. 2022). Activation of the receptor leads to multiple trans-autophosphorylation events of several tyrosine residues in the tyrosine kinase domain (residues Y577, Y579, Y647, Y648 and Y760 in FGFR3) (Ornitz and Itoh 2015). The phosphorylated tyrosines are docking sites for intracellular downstream signaling proteins containing Src homology-2 (SH2) and phosphotyrosine-binding (PTB) domains. Specifically, activated FGFRs phosphorylate and activate the membrane-anchored adaptor protein constitutively associated with the receptor, FGFR substrate 2α (FRS2α), which then binds the growth factor receptor-bound 2 (GRB2) and Src homology-2-containing protein tyrosine phosphatase 2 (SHP2). GRB2 then recruits Son of Sevenless (SOS) and GRB2-associated-binding protein 1 (GAB1). The recruitment of SOS leads to the activation of the RAS - mitogen-activated protein kinase (MAPK) and GAB1 signals downstream to phosphoinositide 3-kinase (PI3K) - protein kinase B (PKB, also known as AKT or PI3K-AKT) pathway. FGFRs also activate a different pathway starting with phosphorylation of phospholipase C *γ* (PLC*γ*), which catalyzes the hydrolysis of phosphatidylinositol 4,5-bisphosphate to inositol triphosphate (IP_3_) and diacylglycerol (DAG). DAG activates protein kinase C (PKC); whereas, IP_3_ leads to an increase of the intracellular calcium ion levels (Ornitz and Itoh 2015).

In addition, FGFRs are able to interact with multiple ligands of the fgf family inducing different levels of activation, which adds yet another layer of complexity to FGFR signaling (Karl et al. 2020; Trenker and Jura 2020). In particular, the splice form FGFR3c used in this study is mainly activated by fgf1, fgf2, fgf8 and fgf9 (Ornitz et al. 1996; Zhang et al. 2006). Of these ligands, fgf1 and fgf2 were reported to show differences in inducing kinase phosphorylation with fgf2 having a stronger activating effect when added at saturating conditions (Sarabipour and Hristova 2016). The resulting outcome of the receptor’s signaling cascade is not only determined by the interplay of different components of the signaling pathways, but also by the number of phosphorylated docking sites in the tyrosine kinase domain, time of activation and inactivation, etc. (Lemmon and Schlessinger 2010). Once FGFR activation has taken place, the signaling cascade needs to be silenced. One common mechanism for signal downregulation is endocytosis that removes the receptor from the plasma membrane. Internalized receptors can be recycled back to the cell surface, rendering them unavailable only for a limited time, or, alternatively, they are permanently eliminated by lysosomal degradation after internalization (Neben et al. 2019). Receptor inactivation is initiated by E3 ubiquitin-protein ligase CBL that forms a complex with phosphorylated FRS2α and GRB2 leading to the ubiquitination of FGFRs and FRS2, which is the signal for receptor internalization (Ornitz and Itoh 2015; Neben et al. 2019).

Furthermore, single point mutations that cause missense amino acid substitutions have been reported to increase the receptor’s activity also in the absence of ligand, and have been defined as gain-of function mutations with profound phenotypic effects causing multiple genetic disorders even in the heterozygous state (reviewed in (Arnheim and Calabrese 2009; Goriely and Wilkie 2012; Arnheim and Calabrese 2016; Tiemann-Boege et al. 2021)). Several missense variants of FGFR3 have been documented in genetic studies of patients with phenotypic disorders such as achondroplasia, thanatophoric dysplasia I and II, Muenke syndrome, hypochondroplasia, among others reported in the Human Gene Mutation database (HGMD) and also in tumors listed in the Catalogue Of Somatic Mutations In Cancer (COSMIC) (Tate et al. 2019). Furthermore, FGFR3 has been categorized as a cancer driver gene involved in cell survival with a 99% oncogene score (van Rhijn et al. 2002a; van Rhijn et al. 2002b; Vogelstein et al. 2013). Over 7% of the sequenced tumors were positive for FGFR3 mutants, some of them affecting the same codon resulting in different amino acid substitutions (cancer.sanger.ac.uk, (Tate et al. 2019)), further highlighting the proliferative properties of FGFR3 mutations. Given that FGFR3 is mainly expressed in the cartilage, brain, lung and the spinal cord (Avivi et al. 1991; Peters et al. 1993), it appears to be most important in the regulation of cartilage growth, being a physiologic negative regulator of chondrocyte proliferation, thus restricting skeletal growth (Foldynova-Trantirkova et al. 2012; Ornitz and Marie 2015). In addition, signaling is tightly regulated by interactions between epithelial and mesenchymal cells during organogenesis, as well as the initiation and proximal-distal growth of the limb bud (Ornitz and Marie 2015). Thus, FGFR3 mutations are mainly associated with chondrodysplasia syndromes (Turner and Grose 2010; Foldynova-Trantirkova et al. 2012; Ornitz and Itoh 2015).

In spite of the strong phenotypic consequences of FGFR3 mutations, not much is known how substitutions affect the receptor’s signaling. Only a few mutations have been validated with experimental data in terms of the effect of the mutation. These include functional studies that experimentally examined the effect of the mutant protein on receptor activation on its signaling (Bellus et al. 2000b; Krejci et al. 2008; He et al. 2012; Sarabipour and Hristova 2016). The potential deleteriousness (both gain-or loss-of-function modifications) of some mutations is also predicted by *in silico* analysis such as CADD or SIFT scores or by merging information from multiple component methods with some experimental data (Ng and Henikoff 2003; Kircher et al. 2014). However, information on the effect of mutations on the protein’s function is still scarce.

For this reason, the purpose of this work was to implement an assay to determine the kinase activity of FGFR3 directly in living cells that can be further used in the analysis of mutants. Currently, the main method to quantify the activation of FGFRs is Western Blotting, which determines the phosphorylation state of tyrosines in the adaptor protein docking sites and compares the signal of the phosphorylated protein to the signal intensity of the pan-protein (Naski et al. 1996; Bellus et al. 2000b; Bonaventure et al. 2007; Gibbs and Legeai-Mallet 2007) or the *in vitro* kinase assay, which uses immunoprecipitated cell lysates that are incubated with ^32^P-ATP (Bellus et al. 2000b). These methods are labor intensive and have other potential pitfalls like poor antibody specificity and electrophoretic resolution, as reviewed in (Ghosh et al. 2014). Moreover, they measure the protein in whole-cell lysates. Particularly for cells artificially overexpressing FGFR3, this might, however, not represent receptor activity at the cell surface, since a fraction of the protein may be still located in intracellular compartments before post-translational modifications (e.g. glycosylation) and translocation to the cell surface take place (De Matteis and Luini 2008). This can be observed in Western Blots as a double-banding pattern formed by the mature receptor that has a different molecular weight than the intracellular fraction (Lievens et al. 2004; Bonaventure et al. 2007; Gibbs and Legeai-Mallet 2007).

In view of the limitations presented by methodologies relying on whole-cell lysates, we here chose to employ a protein micropatterning approach (Schwarzenbacher et al. 2008; Lanzerstorfer et al. 2014; Schutz et al. 2017) to determine the kinase activity of FGFR3 directly in living cells. In this assay, cells are grown on micro-structured surfaces featuring regular arrays of antibodies against the protein of interest (“bait”) with the recruitment of a fluorescently labelled interaction partner (“prey”) being monitored via fluorescence microscopy. The interaction strength between the two proteins can then be quantified via the signal contrast between the prey signal intensity within and outside of the bait regions (Figure 1a) (Schwarzenbacher et al. 2008; Schutz et al. 2017). In the present study, FGFR3 served as prey, and the co-localization of the downstream adaptor protein GRB2 was taken as a measure for the kinase activity of FGFR3 upon stimulation. This experimental design allows to specifically detect only signaling events of the mature protein at the plasma membrane and has been used before to study RTK signaling hubs of the epidermal growth factor receptor (Lanzerstorfer et al. 2014) and kinase recruitment to the T-cell receptor signaling complex (Motsch et al. 2019).

**Figure 1:**
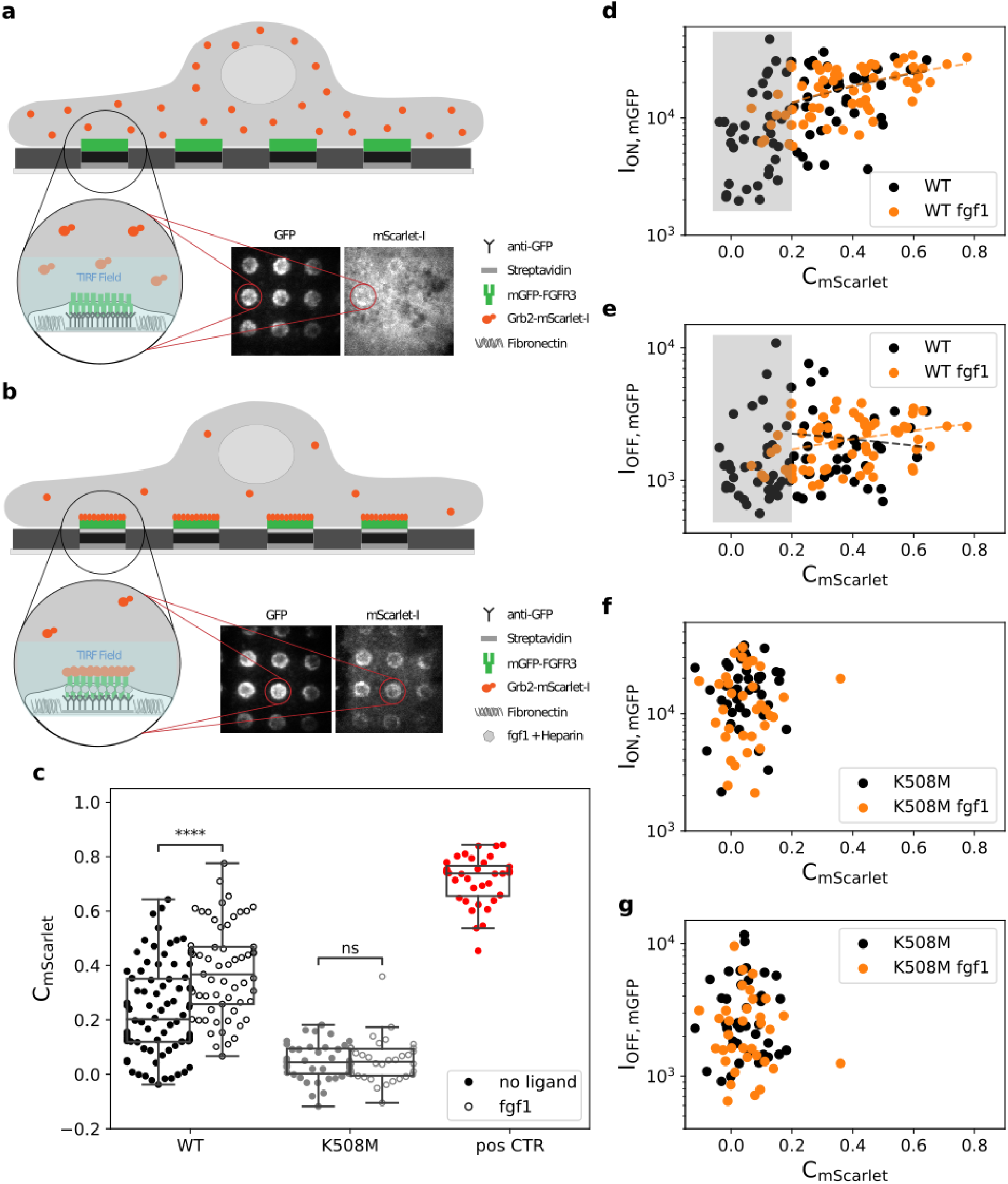
Experimental design and proof-of-principle. **(a**,**b)** Antibody patterns are used to enrich and immobilize mGFP-FGFR3 at specific sites (“ON” regions) in the plasma membrane of HeLa cells, leaving other regions depleted of mGFP-FGFR3 (“OFF”). Co-localization of the adaptor protein GRB2-mScarlet to mGFP-FGFR3 patterns reports on the activation state of FGFR3, with no or little co-patterning observable in the non-activated state (a) and a high degree of co-patterning for the activated receptor after addition of the ligand fgf1 (b). TIR illumination is used to specifically detect membrane-proximal protein. **(c)** The fluorescence contrast of GRB2-mScarlet (C_mScarlet_) relates the fluorescence intensity within ON (I_ON,mGFP_) and OFF (I_OFF,mGFP_) areas of FGFR3-enriched regions and serves to quantify the extent of co-localization. Each dot represents one cell. C_mScarlet_ data for the wildtype receptor (WT), a kinase-dead mutant (K508M) and a mGFP-FGFR3-mScarlet fusion protein as positive control is shown (p-value annotation legend: *= 0.01 ≤ p ≤ 0.05; **= 0.001 ≤ p ≤ 0.01; ***= 0.0001 ≤ p ≤ 0.001; ****= p ≤ 0.0001). **(d**,**e)** Correlation between the receptor’s intensity in ON (d) and OFF (e) regions and the GRB2-mScarlet contrast for the WT receptor. Data in the absence (black) and presence (orange) of fgf1 is shown. The grey box indicates the cell population with C_mScarlet_ < 0.2, which likely represents non-activated cells. **(f**,**g)** Correlation between GRB2-mScarlet contrast and mGFP-FGFR3 intensity in ON (f) and OFF (g) regions for K508M. All correlation coefficients can be found in Supplementary Table S1

Using this micropatterning approach, we were able to quantify the activation of wild type FGFR3, as well as four different pathogenic mutants associated with congenital disorders. Specifically, we focused on two mutations in the transmembrane domain and flanking regions (G380R and Y373C) reported to increase receptor dimerization (He et al. 2011) and sustained ERK activation (Krejci et al. 2008), and two mutations in the kinase domain activation loop (K650Q and K650E). These variants cause different skeletal dysplasias, the latter one being quite severe and embryonic lethal: K650Q (c.1948A>C), hypochondroplasia (HCH) (Bellus et al. 2000b; Leroy et al. 2007); G380R (c.1138G>A), achondroplasia (ACH) (Rousseau et al. 1994; Shiang et al. 1994; Bellus et al. 1995); Y373C (c.1118A>G) thanatophoric dysplasia I (Rousseau et al. 1996); K650E (c.1948A>G) thanatophoric dysplasia type II (TDII) (Tavormina et al. 1995). It has been reported that K650E causes a higher basal level of tyrosine phosphorylation than the G380R mutation, thus providing a possible biochemical explanation why the phenotype of TDII is more severe than that of ACH (Naski et al. 1996; Bellus et al. 2000a; Bellus et al. 2000b). Our results, however, do not corroborate this hypothesis.

## Results

### Characterization of the experimental system

In order to measure FGFR3 activation, HeLa cells expressing mGFP-FGFR3 and GRB2-mScarlet were seeded onto micro-structured surfaces featuring regular patterns of a monoclonal antibody against mGFP (Schutz et al. 2017; Fülöp et al. 2018). The mGFP-FGFR3 at the plasma membrane was enriched and immobilized according to these micropatterns, with the fluorescence intensity reporting on the extent of mGFP-FGFR3 enrichment (I_ON,mGFP_), leaving other regions depleted of mGFP-FGFR3 (yielding I_OFF,mGFP_). The co-recruitment of GRB2-mScarlet to mGFP-FGFR3 patterns was imaged using total internal reflection fluorescence (TIRF) microscopy (Figure 1a,b) with the fluorescence contrast, 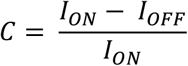serving as a measure for the activation.

First, we determined the level of GRB2-mScarlet recruitment to the mGFP-FGFR3 wild type (WT) construct in the non-liganded state. We observed a pronounced pattern formation in the mGFP channel, reported here as the mean fluorescence contrast for mGFP (mean C_mGFP_ =0.86±0.01, Table 1). While ∼50% of the cells showed a rather uniform distribution of GRB2-mScarlet, the other half exhibited a low to moderate fluorescent contrast for GRB2-mScarlet, yielding a mean C_mScarlet_ = 0.23±0.02 when considering the total number of cells (Figure 1c and Table 1). However, when activating the cells expressing the WT receptor with 50 ng/mL of ligand fgf1 (Ornitz et al. 1996; Ornitz and Itoh 2015) and 1 µg/mL of the co-factor heparin (Schlessinger et al. 2000), we observed a significant increase in GRB2-mScarlet contrast (mean C_mScarlet_ = 0.38±0.02; Figure 1c). The mGFP-FGFR3 contrast stayed within the same range regardless of the ligand addition (mean C_mGFP_ =0.89±0.01 with fgf1, Supplementary Table S2), ruling out that the increase in C_mScarlet_ was a consequence of an increase of C_mGFP_.

**Table 1:** Statistical parameters for all FGFR3 variants tested without ligand. The second column indicates the position of missense mutation in the coding sequence of FGFR3 (CDS) introduced in the expression plasmid. The p-value represents the pairwise comparison of the respective C_mScarlet_ to the non-activated WT. For the complete data set also with the addition of the ligands fgf1 and fgf2 see Supplementary Table S2.

| Variant | Variant (CDS) | Cells | $C_{mScarlet}$ Mean ± SE | p-value | $C_{mGFP}$ Mean ± SE |
| --- | --- | --- | --- | --- | --- |
| K508M | c.1523A>T | 40 | $0.05 \pm 0.01$ | $1.313e-09$ | $0.82 \pm 0.01$ |
| pos. CTR | none | 36 | $0.71 \pm 0.02$ | $1.377e-30$ | $0.93 \pm 0$ |
| WT | none | 76 | $0.23 \pm 0.02$ | n.a. | $0.86 \pm 0.01$ |
| G380R | c.1138G>A | 37 | $0.43 \pm 0.03$ | $2.750e-08$ | $0.92 \pm 0.01$ |
| Y373C | c.1118A>G | 34 | $0.51 \pm 0.03$ | $1.926e-12$ | $0.91 \pm 0.01$ |
| K650Q | c.1948A>C | 35 | $0.54 \pm 0.03$ | $2.780e-15$ | $0.91 \pm 0.01$ |
| K650E | c.1948A>G | 39 | $0.32 \pm 0.03$ | $1.007e-02$ | $0.87 \pm 0.01$ |

As a negative control, we introduced a mutation (K508M) that inhibits the trans-autophosphorylation of the FGFR3 kinase domain and thus recruitment of GRB2, which almost completely abolished GRB2-mScarlet colocalization (mean C_mScarlet_ =0.05±0.01) (Figure 1c). Ligand addition did not induce GRB2 recruitment in cells expressing the kinase-dead mutant K508M (mean C_mScarlet_=0.05±0.01). We also designed a mGFP-FGFR3 construct featuring a C-terminal mScarlet domain (mGFP-FGFR3wt-mScarlet) as a positive control for maximum co-localization, which yielded a mean C_mScarlet_ = 0.71±0.02.

Note that C_mScarlet_ values in the absence, as well as in the presence of fgf1 showed a large variability between individual cells. To ascertain that this variation in C_mScarlet_ was independent of cellular expression levels of the adaptor protein GRB2-mScarlet, we compared it to the adaptor fluorescence intensity ON (I_ON,mScarlet_) or OFF the patterned areas (I_OFF,mScarlet_). We did not find a correlation between C_mScarlet_ and I_ON,mScarlet,_ suggesting that the extent of recruitment of GRB2 to FGFR3 (expressed as C_mScarlet_) is not dependent on differences in expression levels of the transfected cells, but indeed reports on the extent of receptor activation (Supplementary Figure S1). I_OFF,mScarlet_, decreases with C_mScarlet_ as expected, since OFF areas are depleted of GRB2 as it is recruited to the receptor.

### GRB2 colocalization correlates with FGFR3 activation

We next tested whether the contrast of the adaptor protein GRB2 was affected by the expression levels of the receptor. For this purpose, we examined a series of correlations of C_mScarlet_ with the density of FGFR3 in ON (I_ON,mGFP_) and OFF (I_OFF,mGFP_) regions for without fgf1 addition (Figure 1d). Here, we observed the following: i) A cell population (appr. 50%) with C_mScarlet_ > 0.2 showing a weak positive correlation between the GRB2 contrast and ON regions (Spearman’s correlation coefficient between C_mScarlet_ and I_ON,mGFP_ of r = 0.34, significance level 0.04; Figure 1d-e, Supplementary Table S1). ii) A second population of cells of appr. 50% with C_mScarlet_≤0.2 (grey area in Figure 1d-e). This population was reduced when adding the ligand fgf1 (Figure 1d).

We suspected the cell population with contrast levels above 0.2 to represent activated cells. Indeed, for the kinase-dead mutant (K508M) we only observed the population with C_mScarlet_ ≤0.2, likely representing the non-activated cell population (Figure 1f and 1g). For this reason, all further correlation analyses were based on the cell population with high contrast values of C_mScarlet_ > 0.2. Interestingly, even in the absence of ligand a substantial number of cells (∼50%) showed a basal recruitment of GRB2 to the non-ligated WT receptor, possibly due to the trans-autophosphorylation between FGFR3s in the crowded environment inside the micropatterns. As expected though, GRB2 recruitment further increased with the addition of ligand. Note also the increase in the correlation between C_mScarlet_ and I_ON,mGFP_ from r = 0.34 to r = 0.50 with the addition of fgf1 for the cells C_mScarlet_ > 0.2 indicating that the signal we measured was mainly driven by the activation of the receptor (Figure 1d and Supplementary Table S1).

Next, we compared C_mScarlet_ with I_OFF,mGFP_ as an indicator for FGFR3 expression levels (Figure 1e). We did not observe a correlation neither in the absence nor the presence of fgf1 (Supplementary Table S1). This suggests that GRB2 recruitment is rather a consequence of FGFR3 activation within patterns and not just of different FGFR3 expression levels *per se*.

### Effect of disease-relevant mutations on FGFR3 activation

After having established that our experimental set-up is sensitive enough to detect differences in GRB2 recruitment for the non-liganded and fgf1-stimulated WT receptor, we next quantified the FGFR3 activation state for four selected FGFR3 mutations (K650Q, G380R, Y373C, and K650E) described to lead to increased FGFR3 signaling (Figure 2a). Each of these variants causes a different disorder with the latter being very severe and embryonic lethal: HCH, ACH, TDI and TDII. In our experiments, all four variants exhibited a significantly higher mean C_mScarlet_ than the WT receptor without the addition of ligand, indicating the promiscuous recruitment of the adaptor protein GRB2 to the receptor without activation by the ligand (Figure 2b, Table 1). Yet, all variants exhibited mGFP-FGFR3 patterns of similar contrast values (mean C_mGFP_ = ∼0.9), which did not change after fgf1 addition (Table 1, Supplementary Table S2).

**Figure 2:**
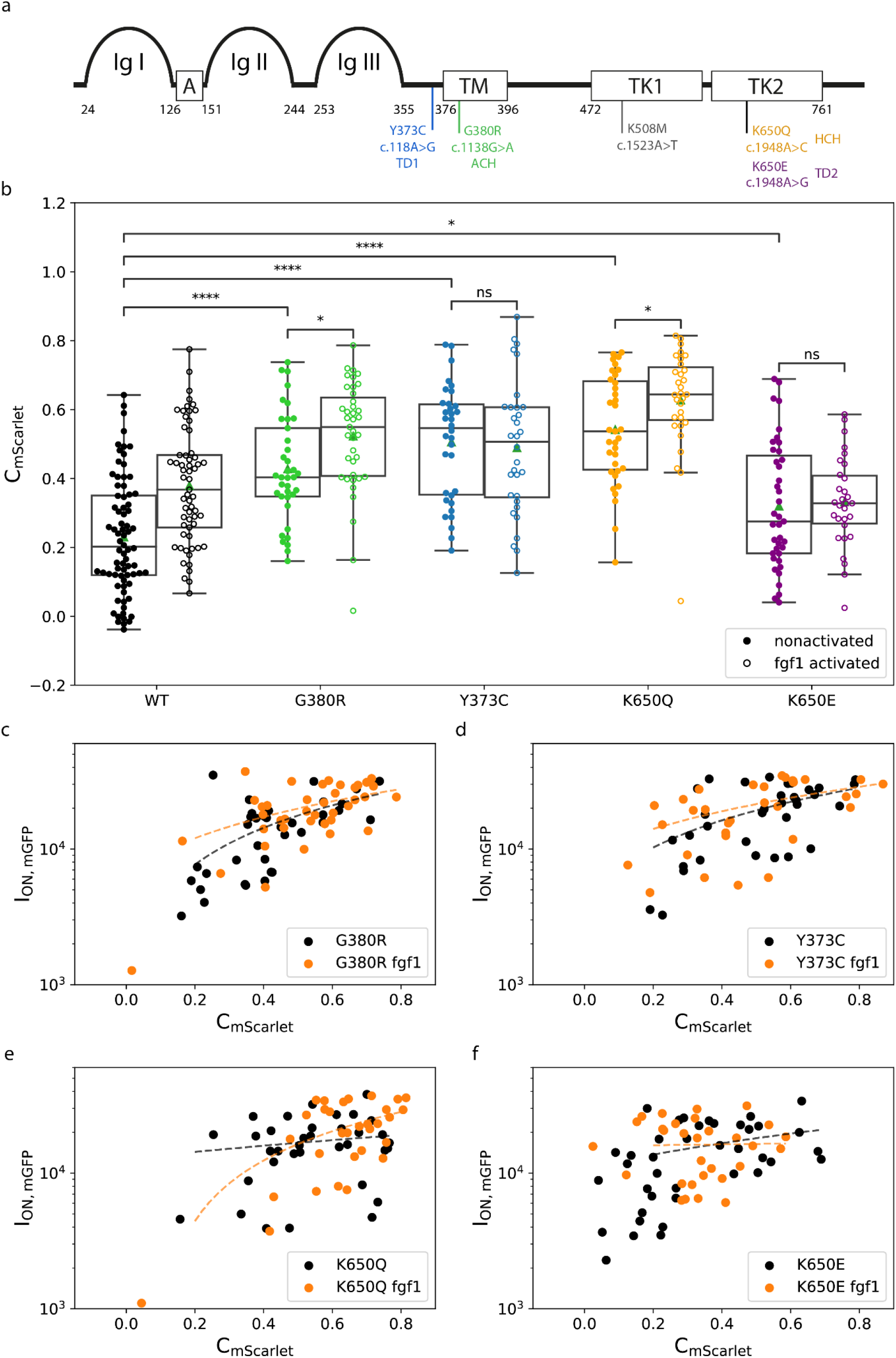
Effect of FGFR3 mutations on receptor activation levels. **(a)** Schematic representation of the FGFR3 receptor and the different tested mutations: Ig-like domains (Ig I-Ig III); acidic box (A), transmembrane domain (TM) and an intracellular split tyrosine kinase domain (TK1 and TK2). Numbers indicate the amino acid position of the respective domains. Mutations are indicated at their approximate location in the protein domains with their respective amino acid and nucleotide substitutions and associated congenital disorder. **(b)** Comparison of the adaptor contrast (C_mScarlet_) determined for the WT and mutant forms of FGFR3 in the absence and presence of fgf1. **(c-f)** Correlation between the receptor intensity in ON areas (I_ON,mGFP_) and the adaptor contrast (C_mScarlet_) for the mutant receptors (Spearman correlation coefficients (r) for each mutant are listed in Supplementary Table S1). The p-value annotation legend is *= 0.01 ≤ p ≤ 0.05; **= 0.001 ≤ p ≤ 0.01; ***= 0.0001 ≤ p ≤ 0.001; ****= p ≤ 0.0001. A full list can be found in Supplementary Table S3.

**Figure 2:**
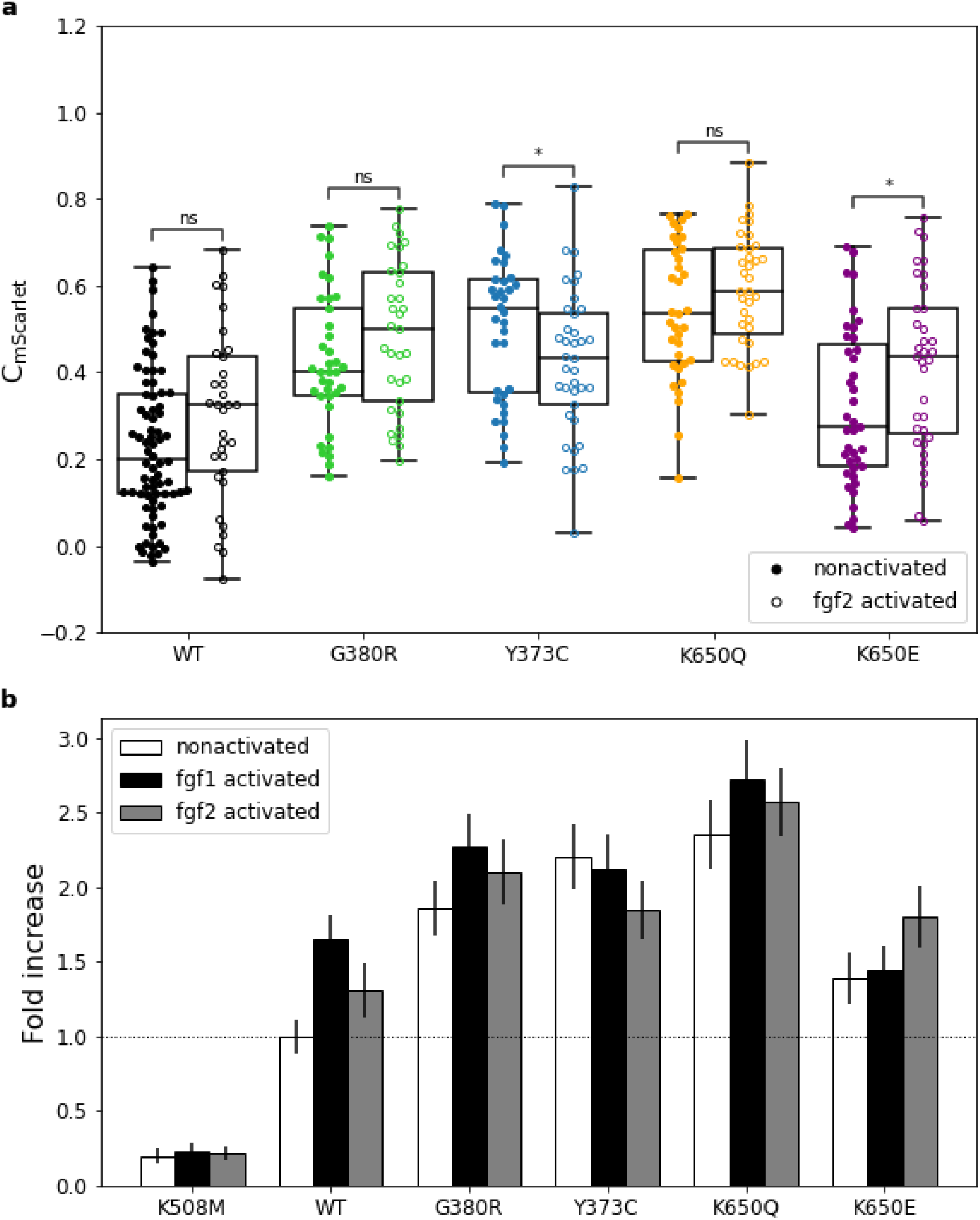
Effect of the ligands fgf1 and fgf2 on receptor activation. **(a)** Comparison of GRB2 contrast (C_mScarlet_) determined for the WT and mutant forms of FGFR3 in the absence and presence of fgf2. The p-value annotation legend is *: 0.01 ≤ p ≤ 0.05; a full list can be found in Supplementary Table S3 **(b)** Normalization of mean C_mScarlet_ values to the non-activated WT FGFR3. Data are shown as mean ± standard error.

The transmembrane domain mutation G380R, which has been suggested to increase the propensity of the receptor to dimerize (He et al. 2010; He et al. 2011; Placone and Hristova 2012), showed an elevated activation level with more GRB2 co-localizing to the receptor (mean C_mScarlet_ = 0.43 ± 0.03) than the WT (Table 1; Figure 2b). Upon addition of fgf1, we observed an increase of this co-localization with a mean C_mScarlet_ = 0.52 ± 0.03 (Figure 2b; Supplementary Table S2) and a shift of the cell population from C_mScarlet_ > 0.2 to C_mScarlet_ > 0.4 (Figure 2c).

Similarly, located in the extracellular juxtamembrane region, the Y373C mutation is hypothesized to induce receptor dimerization via disulfide bond formation causing the constitutive activation of the receptor (d’Avis et al. 1998; Bernard-Pierrot et al. 2006; Bonaventure et al. 2007). Cells expressing the Y373C receptor showed an intrinsic receptor activation with a high GRB2 colocalization without the addition of the ligand (mean C_mScarlet_ = 0.51 ± 0.03) as seen in Table 1 and Figure 2b. Compared to the G380R mutant, this mutant did not have a population of non-activated cells with C_mScarlet_ < 0.2 (Figure 2d). Consistently, addition of fgf1 did not increase the activation of the receptor. Similar to the WT, for both mutants we also observed a positive correlation between the density of FGFR3 on the pattern (I_ON,mGFP_) and C_mScarlet_ without and with the addition of the fgf1 ligand (Figure 2c and d). The Spearman correlation coefficient r was 0.57 for G380R vs. 0.52 for Y373C (Supplementary Table S1), which was higher than for the WT (r = 0.34)

The mutations K650Q and K650E affect the same lysine in the kinase domain leading to strong (K650E; (Webster et al. 1996; Huang et al. 2013)) or moderate (K650Q; (Bellus et al. 2000b)) constitutive receptor activation. Similar to our observations for the G380R variant, K650Q showed elevated FGFR3 activity already in the absence of ligand, which was further increased by fgf1 yielding mean contrast values of 0.54± 0.03 and 0.63 ± 0.03, respectively (Table 1). For this variant, we did not observe a population of non-activated cells, neither in the absence nor the presence of ligand, (Figure 2e). A weak positive correlation between C_mScarlet_ and I_ON,mGFP_ (Spearman’s correlation coefficient r = 0.42) was only found in the presence of ligand (Supplementary Table S1).

Finally, the K650E mutation showed the lowest recruitment of GRB2 to FGFR3 (C_mScarlet_ = 0.32 ± 0.03), in spite of previous studies having reported the K650E mutation as one of the highest activating versions of FGFR3 (Naski et al. 1996; Bellus et al. 2000b). While fgf1 addition did decrease the fraction of non-activated cells (Figure 2f), GRB2 recruitment was still low compared to the WT and the other variants (Figure 2b, Table 1). Further, for K650E we did not observe a correlation between receptor density ON patterns (I_ON,mGFP_) and contrast C_mScarlet_, neither in the absence nor the presence of fgf1 (Supplementary Table S1).

In order to rule out any effect from differences in FGFR3 surface expression, we also tested for correlations between C_mScarlet_ and the receptor density OFF patterns (I_OFF,mGFP_) for all of the analyzed mutants (Supplemental Table S1, Supplemental Figure S2), but did not find any significant correlations neither in the presence nor the absence of fgf1.

### Effect of fgf2 on FGFR3 activation

We further analyzed the different mutations using fgf2 (25 ng/mL with 1 µg/mL heparin), an alternative ligand also reported to activate the isoform IIIc of FGFR3 (Ornitz et al. 1996). The concentration of fgf2 was chosen as reported in the literature (Balek et al. 2018). Fgf2 increased the activation of the wild type receptor, albeit with a less pronounced effect than fgf1 (Figure 3a, C_mScarlet_: nonactivated 0.23± 0.02 vs fgf2 0.3 ± 0.03). While we did not detect a significant increase of C_mScarlet_ upon addition of fgf2 for the mutants G380R and K650Q. The K650E mutant did show a modest increase in GRB2-mScarlet recruitment (see Supplementary Table S2). Conversely, Y373C showed a modest decrease in GRB2-mScarlet recruitment upon fgf2 addition compared to the non-ligated state. In short, fgf2 has a milder effect compared to fgf1 but this could be also explained by the lower concentration used for fgf2 compared to fgf1 (25ng/ml vs 50ng/ml).

In accordance with our fgf1 data, we also tested the fgf2-activated cells for correlations of C_mScarlet_ to either I_ON,mGFP_ or I_OFF,mGFP_. Similar to the fgf1 data, non-activated cells (C_mScarlet_ <0.2) almost completely disappeared after addition of fgf2 for the WT and the mutants G380R and K650Q (Supplemental Table S1 and Supplemental Figure S3) even though the mean C_mScarlet_ with fgf2 was slightly lower compared to fgf1 (Supplemental Table S2). For K650Q we also found a positive correlation between C_mScarlet_ and I_ON,mGFP_ after addition of fgf2, as well as for Y373C and K650E (Supplemental Table S1). Similar to the wild type or the other mutants there was no correlation, except for G380R that shows a negative correlation between C_mScarlet_ and I_OFF,mGFP_ (Supplemental Table S1).

In order to better visualize the differences in receptor activation among the different mutants, as well as their respective behavior in response to ligand addition, we normalized the mean C_mScarlet_ to the non-ligated WT receptor (Figure 3b). It can be observed that the GRB2 recruitment to the WT receptor in the absence of ligand is significantly increased (∼5-fold) compared to the negative control K508M. The addition of fgf1 to the WT leads to a ∼1.5-fold increase in C_mScarlet_ compared to the non-stimulated WT; in contrast, the fgf2 ligand only leads to a ∼1.3-fold increase. All the mutants exhibit at least 1.5-fold increased GRB2 recruitment in the unligated state, with the K650Q mutation showing the highest increase of about 2.5-fold. The overall highest signaling can be observed in the fgf1 activated K650Q mutant which reaches almost 3-times the level of the non-induced WT.

## Discussion

### Quantification of FGFR3 signaling in live cells on micropatterned surfaces

In this work, we used a protein micropatterning approach (Lanzerstorfer et al. 2014; Motsch et al. 2019) to quantify FGFR3 activity in living cells. By monitoring the recruitment of a downstream signaling molecule (in our case GRB2) to FGFR3 at the plasma membrane (Lanzerstorfer et al. 2014; Motsch et al. 2019), we directly captured the activity of the mature receptor at the cell surface, in contrast to the more commonly used methods that analyse whole cell lysates (e.g. Western Blotting). This aspect is important since in this case endocytic vesicles and/or partially processed receptor in the ER or Golgi apparatus contribute to the detected signal, thus distorting the results (Lievens et al. 2004; Bonaventure et al. 2007; Gibbs and Legeai-Mallet 2007). We found that all four FGFR mutations tested in our study yielded in a promiscuously active surface receptor with higher levels of GRB2 recruitment than the wildtype even in the absence of ligand. Further, we quantified the extent of activation in response to two different FGFR3 ligands, fgf1 and fgf2. We observed that ligand addition resulted in an additional increase in GRB2 recruitment for G380R and K650Q, but not for the variants Y373C and K650E.

In our system, the receptor is enriched and immobilized according to specific patterns in the plasma membrane. As expected, we observed an increased recruitment of GRB2 to FGFR3-enriched areas, indicated by an increase of the contrast (mean C_mScarlet_) when adding the ligand. The recruitment was also proportional to the abundance of the receptor in the patterns (I_ON,mGFP_) (Figure 1d and 1e). Interestingly, in a subpopulation of cells we observed already some GRB2 recruitment in the WT without the addition of the ligand. This basal recruitment might be driven by the trans-autophosphorylation of FGFR3s forming dimers or multimers in the crowded environment of the micropatterns. It has been described that a different RTK (EGFR) can form dimers in the absence of ligand leading to ligand-independent activation (Paul and Hristova 2019). Moreover, this RTK has an optimal activity as an oligomer compared to the dimer form (Paul and Hristova 2019). Of the three FGFRs, FGFR3 has the highest intrinsic propensity for dimerization even in the absence of ligand (Sarabipour and Hristova 2016). Further, the receptor activity can be triggered by a higher receptor density, which is a common mechanism to increase the signaling output in cancer cells associated with FGFR upregulation (Turner and Grose 2010). Thus, we hypothesize that the clustering of FGFR3 in our micropatterns enhances the receptor activation in the absence of ligand.

### Transmembrane mutants show promiscuous activation

We showed that four different FGFR3 mutations associated with human congenital disorders pre-activate the signaling of the receptor at the plasma membrane. Specifically, we analyzed the FGFR3 mutations K650Q, G380R, Y373C, and K650E described to lead to increased FGFR3 signaling and different disorders of increasing severity (HCH, ACH, TDI and TDII, respectively). The mutations Y373C and G380R are located next or within the transmembrane domain, respectively. The Y373C variant has been described to lead to the formation of covalently bound dimers, formed by a disulfide bond between the free cysteine residues introduced by the mutation (d’Avis et al. 1998). Similarly, the variant G380R introduces a positively charged residue into the main dimer interface created by interactions between the residues Y379, F384 and F386 (Bocharov et al. 2013). Thus, also this mutation stabilizes FGFR3 dimers already in the absence of ligand, but to a lesser extent than the covalent bond in case of Y373C, and leads to a weak increase in receptor activity (Naski et al. 1996). In accordance with these observations, our study showed i) ligand-independent activation of both mutants in the basal state, with Y373C showing a higher level of activation than G380R (Figure 2b and 3b); and ii) a further increase of activity upon stimulation with fgf1 only for the G380R variant (Figure 2b and 3b). Finally, the higher activation of Y373C compared to G380R provides a plausible explanation of the phenotypic differences of TDI (caused by Y373C) with ACH (caused by G380R), with the former being often more severe.

### Activation behavior of kinase domain mutations

In our study we did not observe a direct correlation between the activation level and the phenotypic severity for all tested mutants, as was proposed in earlier studies for other mutations of FGFR3 (Naski et al. 1996; Bellus et al. 2000a; Bellus et al. 2000b). For the mutations in the kinase domain (K650E and K650Q), the activation behavior of the mutant receptors was more complex. The K650E variant changes the conformation of the tyrosine kinase domain similar to ligand-mediated FGFR3 dimerization and autophosphorylation (Webster et al. 1996; Huang et al. 2013). The K650Q variant has been described to also result in constitutive kinase activation; however, to a lesser degree than the K650E mutant (Bellus et al. 2000b). In contrast, we found that the high basal level of activity of the K650Q variant could be enhanced even further by the addition of fgf1. Pertaining to the phenotypic consequences of the mutation, K650Q is associated with HCH, which is considered a milder form of ACH (Foldynova-Trantirkova et al. 2012), and also milder than the other skeletal dysplasias TDI and TDII. Thus, this particular example (K650Q) with the highest promiscuous activation measured in our system (∼3-fold higher than WT) does not follow the expectation that the severity of the skeletal dysplasia is correlated with the degree of the activation of the mutant receptor.

Another intriguing observation is that, contrary to previously published data, the kinase mutation K650E had the lowest GRB2 recruitment of all four analyzed mutations (Figure 2b and 3b), although it has been described as one of the mutations leading to the highest constitutive receptor activation (Naski et al. 1996; Webster et al. 1996; Huang et al. 2013). A partial, and maybe still unsatisfactory, explanation for these contradictory observations, might be the cellular compartment harboring the mutant protein. Previous studies have shown that mutant forms of the FGFR3 intracellular protein chain are not released and remain inside the cellular export machinery (Lievens et al. 2004; Bonaventure et al. 2007; Gibbs and Legeai-Mallet 2007). While these intra-cellular fractions can still induce basal activation of the downstream ERK proteins (Lievens et al. 2004; Bonaventure et al. 2007; Gibbs and Legeai-Mallet 2007), the biological importance of these immature receptors is still unclear. Studies analyzing the activation in whole cell lysates (Naski et al. 1996; Webster et al. 1996; Bellus et al. 2000b) would capture the receptor in the intracellular compartments and in the cell membrane. Our system specifically reports on signaling events happening at and/or near the plasma membrane and thus responding to ligand induction.

### Fgf1 vs fgf2 ligand activation

In this study we determined differences between the fgf1 and fgf2 ligands for recruiting the adaptor GRB2. Using the micropatterning approach, we observed that fgf1 leads to more GRB2 recruitment to the WT receptor. The addition of fgf2 did elicit a weaker activation (Figure 2b and 3b), which is in line with previous findings showing that fgf1 has a higher affinity for FGFR3 isoform IIIc (also used in our study) and binds it more stably than fgf2. This stable binding results in a longer signaling response and a stronger activation (Chen and Hristova 2011).

A different study from the same lab using a truncated FGFR3 form and highly saturating conditions for both fgf1 and fgf2 (5 µg ml^-1^) observed an opposite effect: fgf1 induced a lower phosphorylation activity than fgf2 (Sarabipour and Hristova 2016). The reported physiologic abundance of the ligands detected in plasma of healthy individuals is approx. 4 ng ml^-1^ for fgf1 (Liang et al. 2018) and at approx. 107 pg ml^-1^ for fgf2 (Yeboah et al. 2007). Although we used ligand concentrations far from these physiological conditions (50 ng/ml fgf1 or 25 ng/ml fgf2), the 2-fold higher concentration of fgf1 than fgf2 might account for the stronger effect we observed. However, note that fgf1 is less stable compared to fgf2 (Buchtova et al. 2015)), so it is difficult to estimate the effective ligand concentration in our system. Further we also used heparin that has also a strong stabilizing effect on fgf1 but a more moderate effect on fgf2 (Buchtova et al. 2015).

In conclusion, the effect of mutations on the activation behavior of FGFR3 is diverse; as reported here, some mutants are still responsive to ligand binding, but others already result in a very high promiscuous receptor activity and do not further increase their activity upon ligand addition (e.g. Y373C compared to G380R), which is of importance for therapies using ligand inhibitors. The method we present here complements existing approaches in that it allows to test these activation differences directly at the plasma membrane in living cells and measure the response to the ligand.

## Methods

### Cloning and plasmids

The aim was to create a fusion construct containing the coding sequence of the human *FGFR3* gene (isoform 1, also known as *FGFR3-IIIc*) with an N-terminal monomeric GFP (mGFP) tag. The sequence of the mGFP contains a mutation that prevents dimerization of the fluorophore (Zacharias et al. 2002). The final fusion-protein was cloned into the pcDNA3.1/Hygro(+) vector (ThermoFisher Scientific), suitable for expression in mammalian cells. For proper intracellular transport and secretion of the fusion protein, the signal peptide (SP) was cloned N-terminal of the mGFP, followed by a 5xGGS-linker that serves as a spacer between the GFP and the other domains of FGFR3 (extracellular domain, EC; transmembrane domain, TM; intracellular domain, IC); FGFR3(SP)-mGFP-5xGGS-FGFR3(EC-TM-IC). To ensure efficient translation initiation, the strong Kozak sequence “GCCACC” (Kozak 1987) was cloned upstream of the “ATG”-start codon. Cloning was performed by restriction enzyme digest and ligation in two steps. In step 1, the FGFR3 domains without the signal peptide (all primer sequences used in this study are shown in Supplementary Table S4) were cloned into the empty pcDNA3.1/Hygro(+) vector and in step 2 the generic construct FGFR3(SP)-mGFP-5xGGS, which was synthesized and cloned by BioCat (the sequence is shown in Supplementary Table S5), was cloned upstream of FGFR3. A detailed description of the different steps of the cloning procedure are described in the Supplementary Material and Methods.

In order to introduce the specific single-point mutations studied in this paper, the FGFR3 expression vector was amplified using a high-fidelity polymerase, either Q5 (NEB, USA) or Phusion HS II (ThermoScientific, USA). Specifically designed (back-to-back) primers, where one primer per primerpair carries a 5’-phosphate modification, were used for the amplification, thereby introducing site-directed base changes that create non-synonymous (and in addition several silent) mutations within the *FGFR3* gene. The silent mutations were inserted in close proximity to the actual mutation, to detect potential aerosol contaminations of these plasmids in ultra-sensitive sequencing technologies used in other projects in our laboratory. To avoid potential side effects due to different codon usage, the silent mutations were designed such that codons with a similar usage frequency in humans were chosen. A list of all mutated plasmids generated for this study is shown in Supplementary Table S6. Description of the detailed mutagenesis procedure is given in Supplementary Material and Methods.

To obtain the GRB2-mScarlet plasmid, we carried out PCR to amplify the *mScarlet* sequence from the ITPKA-mScarlet plasmid (Addgene, USA) as well as the *GRB2* sequence from GRB2-YFP (gift from J. Weghuber, FH Wels, Austria) with >15 nt overhangs complementary to adjacent regions on the target plasmid. We then used the Gibson assembly Master Mix (E2611, NEB, USA) following the supplier’s instructions to insert both fragments into a pcDNA3.1 vector.

The positive CTR plasmid was created by PCR amplification of the complete vector containing the mGFP-FGFR3 expression construct using the high-fidelity polymerase Q5 (NEB, USA). Specific primers were designed to delete the stop codon of *FGFR3* and to also add the recognition sequence for the *AgeI* restriction enzyme. In another PCR reaction using Q5, the *mScarlet* was amplified from our GRB2-mScarlet plasmid using primers that also contained the *AgeI* recognition sequence as well as additional bases that are translated into “GS” in order to generate a GGS-linker in between the FGFR3 and the mScarlet. Description of the detailed cloning procedure is given in Supplementary Material and Methods.

### Cells and reagents

HeLa cells (human cervical cancer cells, ACS7005 Sigma-Aldrich, USA) were cultured in DMEM (Dulbecco’s Modified Eagle Medium) medium supplemented with 10 % fetal bovine serum (FBS), 2 mM L-glutamine, 1000 U/ml penicillin-streptomycin (all from Sigma-Aldrich, USA) in a humidified atmosphere at 37 °C and 5 % CO_2_. Cells were co-transfected with one of the pcDNA3.1-mGFP-FGFR3 construct and the pcDNA3.1-GRB2-mScarlet in Opti-MEM I reduced serum media (ThermoScientific, USA) using TurboFect™ Transfection Reagent (ThermoScientific, USA), according to the manufacturer’s protocol. The imaging buffer used for microscopy consisted of HBSS (Sigma-Aldrich, USA) supplemented with 2 % FBS. FGFR3 receptor expressing cells were activated using FGFR ligands fgf1 or fgf2 (bio-techne, USA).

### Surface preparation and activation of patterned cells

Micropatterned surfaces were produced as previously published (Schutz et al. 2017). In brief, polydimethylsiloxane (PDMS) stamps featuring circular pillars with diameters of 3 µm and a center-to-center distance of 6 µm were rinsed with absolute ethanol and dH_2_O and dried in a N_2_ flow. followed by 15 minutes incubation with 50 µg/ml streptavidin (Sigma-Aldrich, USA) in phosphate-buffered saline (PBS, Sigma-Aldrich, USA). After incubation, the polymer stamps were rinsed with dH_2_O, dried in a N_2_ flow, then placed on an epoxy-coated coverslip (Schott, Germany), pressed slightly to ensure good contact between the surfaces and incubated for 60 min at room temperature (RT). After stamp removal, a Secure-Seal hybridization chamber (Grace BioLabs, USA) was placed on top of the streptavidin pattern and filled with 50 µg/ml fibronectin (Sigma-Aldrich, USA) in PBS. After 30 min fibronectin was removed and 10 µg/ml biotinylated anti-GFP (Novus, USA) in PBS with 1 % bovine serum albumin (BSA, Sigma-Aldrich, USA) was added for 30 min, followed by rinsing with PBS. For micropatterning experiments, cells were harvested approximately 24 hours after the transfection using Accutase (Sigma-Aldrich, USA), seeded onto micropatterned surfaces and incubated for 30 minutes at 37°C. Directly before the measurement, the medium was replaced with imaging buffer. After experiments with cells in the non-activated state, the hybridization chamber was filled with imaging buffer supplemented with 1 µM heparin and incubated at 37 °C for 3 min. Then the solution in the chamber was exchanged to imaging buffer with either 50 ng/ml fgf1 or 25 ng/ml fgf2 and the sample was incubated for 15 min at 37 °C.

### Total internal reflection fluorescence (TIRF) microscopy

TIRF microscopy experiments were performed on a home-built system based on a modified inverted microscope (Zeiss Axiovert 200, Germany) equipped with 100x oil-immersion objective (Zeiss Apochromat NA1.46, Germany). mGFP was excited using a 488 nm diode laser (ibeam-smart, Toptica, Germany) and mScarlet was excited using a 561 nm diode laser (Obis, Coherent, USA). Laser lines were overlaid with an OBIS Galaxy beam combiner (Coherent, USA). Emission light was filtered using appropriate filter sets (Chroma, USA) and recorded on an iXon DU 897-DV EM-CCD camera (Andor, Ireland). TIRF illumination was achieved by shifting the excitation beam in parallel to the optical axis with a mirror mounted on a motorized movable table.

### Data evaluation of contrast analysis

Images of patterned cells were analyzed using Fiji (Schindelin et al. 2012). Briefly, selection masks defining “ON” and “OFF” were determined based on the FGFR3-mGFP pattern and applied on the images recorded in the GRB2-mScarlet channel. All “ON” and all “OFF” areas of one cell (usually 4-9) were pooled and the background corrected mean pixel intensity values of “ON” and “OFF” regions, I_ON_ and I_OFF_, were used for further analysis. The contrast value was then determined separately for each cell and color channel via

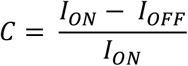

For each experiment, we collected at least 30 individual data points (cells) from at least 3 independent measurements (transfections) were merged for the contrast plots and the corresponding mean values were compared using a one-way-ANOVA. Exact cell numbers are presented in the respective tables. Correlation between GRB2 contrast and GFP ON pattern intensity was determined by Pearson’s correlation coefficient. Statistical analysis and plotting were implemented in Python 3, using the Numpy, SciPy and Pandas packages for general numerical computations, Matplotlib and Seaborn for plotting and Statannot for statistical annotation in plots.

## Data Access

Abbreviations

## Disclosure Declaration

### Ethics approval and consent to participate

Not applicable

### Consent for publication

Not applicable

***No Competing interests***

## Acknowledgements

We would like to thank Pavel Krejci for advice and fruitful comments and general information on FGFR3 activity, and for help in the conceptualization stages of the assay. We would also like to thank Kalina Hristova for the initial WT construct of FGFR3.

## Author Contributions

IH, VB, YS and AY performed experiments and data analysis. GS and ITB conceived the project and provided funding. IH, VB, ES and ITB wrote the manuscript. All authors discussed the data and read and approved the MS.

