## Supplementary Figures for "Quantitation of FGFR3 signaling via GRB2 recruitment on micropatterned surfaces"

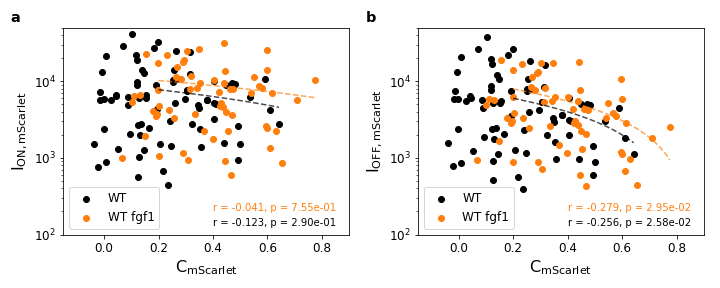
Figure S 1: Correlation analysis of GRB2-mScarlet contrast (C_mScarlet_) and intensity for WT FGFR3 before (black) and after (orange) the addition of fgf1 in (a) ON areas (I_ON,mScarlet_) and (b) OFF areas (I_OFF,mScarlet_). Pearson's correlation coefficient (r) is shown as the dotted line in the respective color.

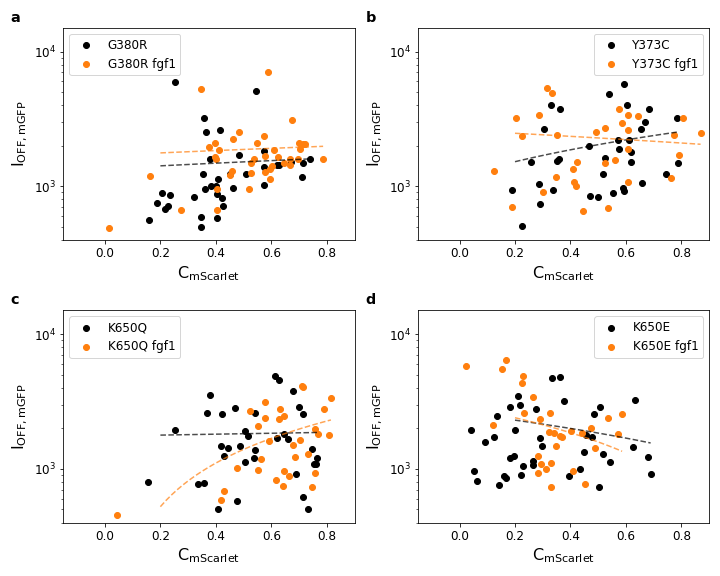

Figure S 2: Correlation between the mGFP-FGFR3 intensity in OFF regions (I_OFF,mGFP_) and GRB2-mScarlet contrast (C_mScarlet_) for G380R (a), Y373C (b), K650Q (c) and K650E (d). Data in the absence (black) and presence (orange) of fgf1 is shown. Pearson's correlation coefficient (r) is shown as the dotted line in the respective color. The values of (r) are reported in Supplemental Table 1.

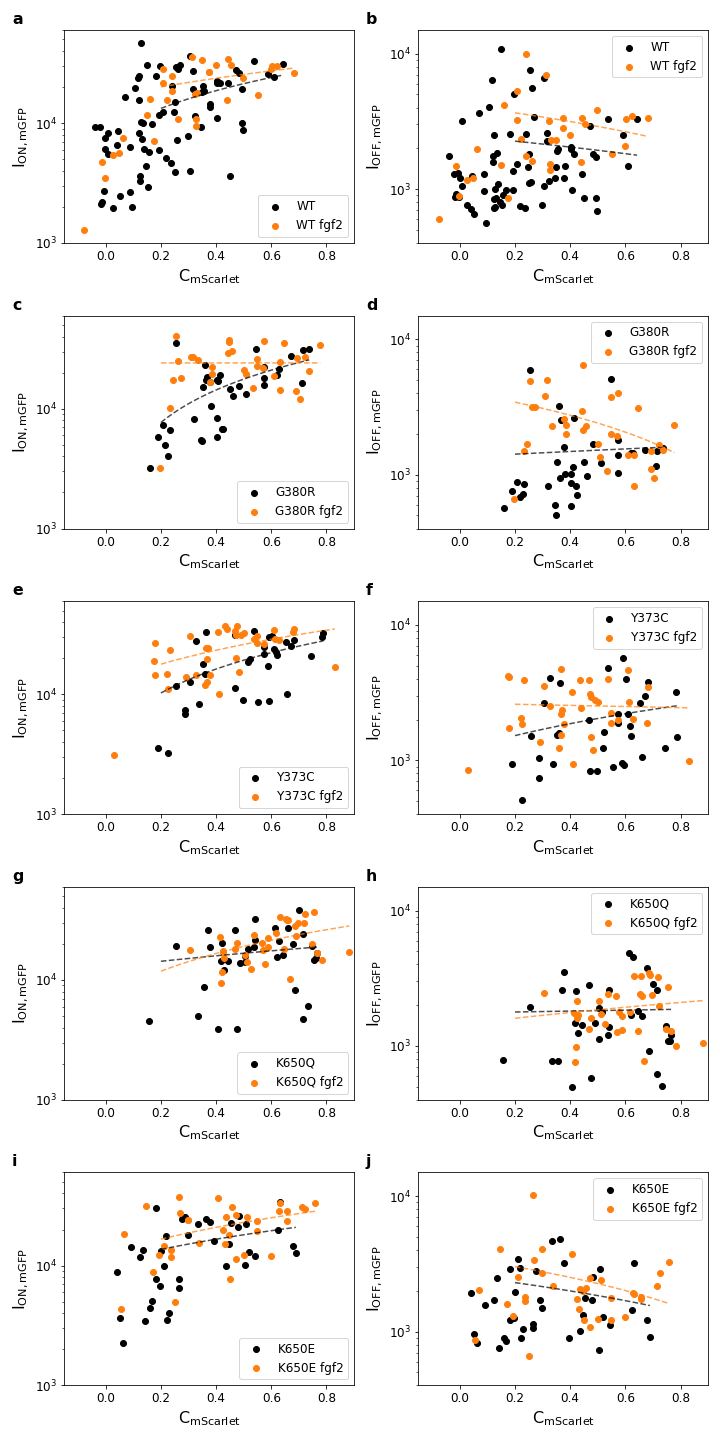

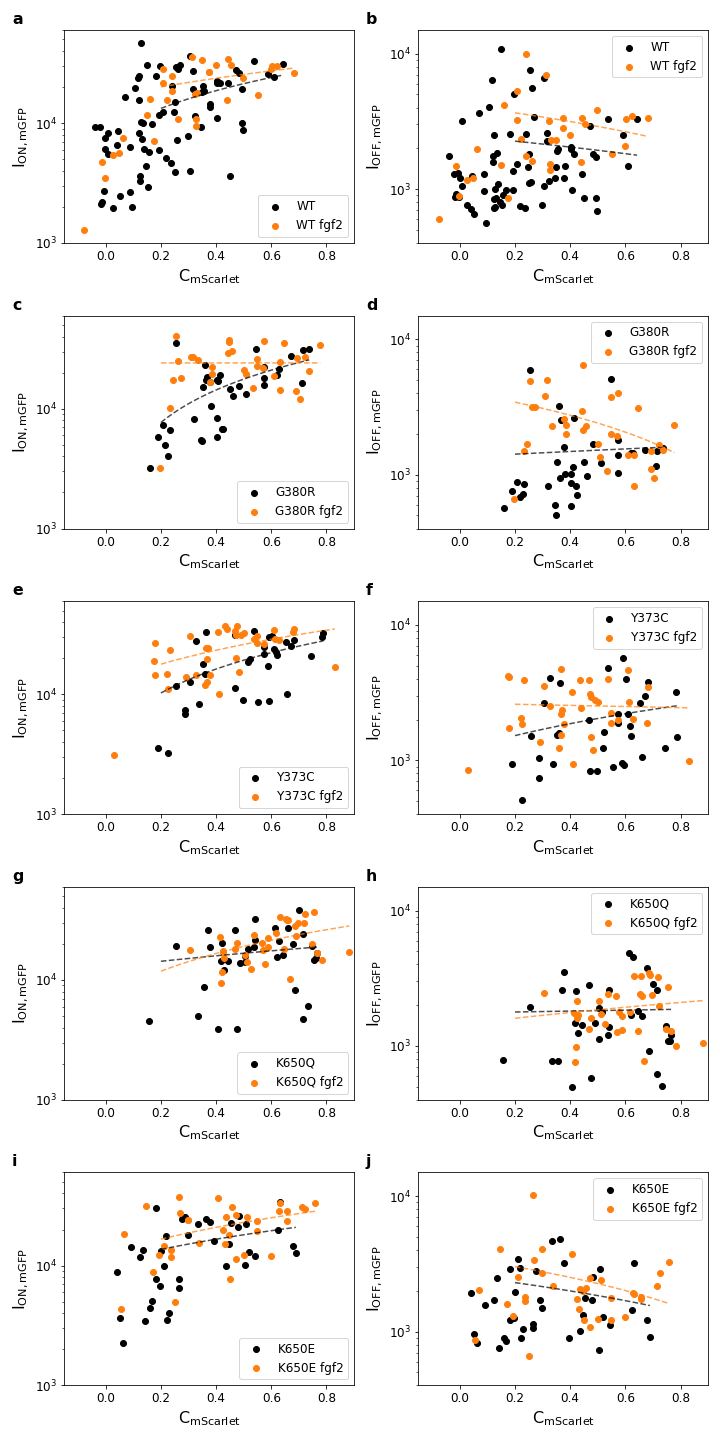

Figure S 3: Correlation between mGFP intensity in ON (I_ON,mGFP_) (left panel; a, c, e, g and i) and OFF regions (I_OFF,mGFP_) (right panel; b, d, f, h and j) and GRB2-mScarlet contrast (C_mScarlet_) for the WT and all the analyzed mutants. Data in the absence (black) and presence (orange) of fgf2 is shown. Pearson's correlation coefficient (r) is shown as the dotted line in the respective color. The values of (r) are reported in Supplemental Table 1.
