## Supplementary Methods for "Quantitation of FGFR3 signaling via GRB2 recruitment on micropatterned surfaces"

**Cloning of mGFP-FGFR3 into pcDNA3.1**

*Step 1: Cloning of FGFR3(EC-TM-IC) into pcDNA3.1/Hygro(+)*

In the beginning, the coding sequence of the extracellular-, transmembrane-, and intracellular domain of the human FGFR3 was amplified from a vector template containing the human *FGFR3-IIIc* gene with a silent mutation at position c.882T>C (p.N294N), (kindly provided by Kalina Hristova, Department of Materials Science and Engineering, Johns Hopkins University, Baltimore, Maryland 21218, USA; “pcDNA3.1-FGFR3-YFP vector”), using the primer set BamHI-FGFR3(noSP)_F and FGFR3(noSP)-XbaI_R (primer sequences are shown in Supplemental Table S4). The template vector was linearized by digestion using 10 units (U) of the restriction enzyme *HindIII*-HF (NEB) in 1x CutSmart buffer using an incubation time of 90 min at 37 °C followed by dephosphorylation using the Antarctic Phosphatase (NEB) in 1x Antarctic Phosphatase Buffer for 15 min at 37 °C, concluded by a heat inactivation of 20 min at 80 °C. Q5-High-Fidelity DNA Polymerase (NEB) was used at 0.3 U/µl in a 25 µl reaction in 1x Q5-reaction buffer supplemented with 1x GC-enhancer, 0.2 mM dNTPs (Biozym) and 0.4 µM of each primer. 1 ng of the linear vector DNA was used as template. The PCR started with an initial heating step of 98 °C for 30 sec, followed by 5 cycles at 98 °C for 10 sec, 65.5 °C for 15 sec, and 72 °C for 70 sec, followed by 20 cycles at 98 °C for 10 sec and 72 °C for 85 sec, concluded by a final elongation step of 3 min at 72 °C in a T100 Thermal Cycler (BioRad). The correct length of the amplicon(s) was assessed via gel electrophoresis and the desired band was excised from the agarose gel using a clean scalpel and purified using the Wizard SV Gel and PCR Clean-Up System (Promega) according to manufacturer’s instructions. The amplicon was eluted in 30 µl nuclease-free water and the DNA concentration was determined using a Nanodrop 2000 instrument (Thermo Scientific).

The purified PCR amplicon was digested with 10 units of the restriction enzymes *BamHI*-HF (NEB) and *XbaI* (NEB) in 1x CutSmart Buffer using an incubation time of 60 min at 37 °C followed by a heat inactivation of 20 min at 80 °C. After the digest, a buffer exchange was performed using the Wizard SV Gel and PCR Clean-Up System (Promega) according to manufacturer’s instructions. The insert-DNA was eluted in 30 µl nuclease-free water and the DNA concentration was determined using a Nanodrop 2000 instrument (Thermo Scientific). The vector DNA (500 ng) was digested with 10 units of the restriction enzymes *BamHI*-HF (NEB) and *XbaI* (NEB) in 1x CutSmart Buffer using an incubation time of 90 min at 37 °C followed by dephosphorylation using the Antarctic Phosphatase (NEB) in 1x Antarctic Phosphatase Buffer for 15 min at 37 °C, concluded by a heat inactivation of 20 min at 80 °C. The desired band was excised from a 1% agarose gel using a clean scalpel and the linearized vector DNA was purified using the Wizard SV Gel and PCR Clean-Up System (Promega) according to manufacturer’s instructions. The vector-DNA was eluted in 30 µl nuclease-free water and the DNA concentration was determined using a Nanodrop 2000 instrument (Thermo Scientific).

Ligation was performed using 50ng of vector-DNA mixed with insert-DNA in a 1:3 vector-to-insert ratio in a 20 µl reaction supplemented with 1x T4 DNA Ligase Reaction Buffer (NEB) and 400 units T4 DNA Ligase (NEB). The reaction was mixed by pipetting and incubated over night at 16 °C. Afterwards, 10 µl of the ligation reaction were transformed into 50 µl of chemically competent *E. coli* XL1 Blue (Agilent) according to manufacturer’s instructions and plated on LB-agar containing 100 µg/mL ampicillin. Screening for positive clones was performed by colony PCR using the primer set FGFR3_pos2343_F and BGH(pcDNA3.1)_R (primer sequences are shown in Supplemental Table S4). OneTaq® (NEB) polymerase was used at 0.025 U/µl in a 20 µl reaction in 1x standard buffer supplemented with 0.2 mM dNTPs (Biozym) and 0.5 µM of each primer. As template, the single colonies were dipped into the PCR master mix. The PCR started with an initial heating step of 94 °C for 5 min, followed by 30 cycles at 94 °C for 15 sec, 57 °C for 20 sec, and 68 °C for 15 sec, concluded by a final elongation step of 5 min at 68 °C. The correct length of the amplicon was assessed via gel electrophoresis and the positive clones were inoculated in 5 mL LB medium containing 100 µg/mL ampicillin to grow an overnight culture, shaking at 37 °C. 4 mL of the culture were harvested by centrifugation and a plasmid Miniprep was performed using the PureYield Plasmid Miniprep System (Promega) according to manufacturer’s instructions. The pure plasmids were sent to sequencing at LGC genomics.

*Step 2: Cloning of FGFR3(SP)-mGFP-5xGGS into FGFR3(EC-TM-IC)_pcDNA3.1/Hygro(+)*

The desired FGFR3(SP)-mGFP-5xGGS fragment (sequence shown in Supplemental Table S5) was purchased from BioCat as an insert in pUC57 vector between the restriction sites *HindIII* and *BamHI*. The pUC57-vector clone containing the verified sequence of the FGFR3(SP)-mGFP-5xGGS fragment was used for restriction enzyme digest to isolate the insert for further cloning. 500 ng vector DNA were digested with 10 units of the restriction enzymes *HindIII*-HF (NEB) and *BamHI*-HF (NEB) in 1x CutSmart Buffer using an incubation time of 60 min at 37 °C followed by a heat inactivation of 20 min at 80 °C. The desired band was excised from a 1% agarose gel using a clean scalpel and the fragment was purified using the Wizard SV Gel and PCR Clean-Up System (Promega) according to manufacturer’s instructions. The insert-DNA was eluted in 30 µl nuclease-free water and the DNA concentration was determined using a Nanodrop 2000 instrument (Thermo Scientific). 500ng of the FGFR3(EC-TM-IC)_pcDNA3.1/Hygro(+) vector (created in step 1) were digested with 10 units of the restriction enzymes *HindIII*-HF (NEB) and *BamHI*-HF (NEB) in 1x CutSmart Buffer using an incubation time of 60 min at 37 °C followed by a heat inactivation of 20 min at 80 °C. The desired band was excised from a 1% agarose gel using a clean scalpel and the linearized vector DNA was purified using the Wizard SV Gel and PCR Clean-Up System (Promega) according to manufacturer’s instructions. The vector-DNA was eluted in 30 µl nuclease-free water and the DNA concentration was determined using a Nanodrop 2000 instrument (Thermo Scientific). The ligation and transformation were carried out as described before. Lastly, for colony PCR the primer set CMV_sequ_F and GFP_pos61_R was used (primer sequences are shown in Supplemental Table S4). OneTaq® (NEB) polymerase was used at 0.025 U/µl in a 20 µl reaction in 1x standard buffer supplemented with 0.2 mM dNTPs (Biozym) and 0.5 µM of each primer. As template, the single colonies were dipped into the PCR master mix. The PCR started with an initial heating step of 94 °C for 5 min, followed by 30 cycles at 94 °C for 15 sec, 57 °C for 20 sec, and 68 °C for 30 sec, concluded by a final elongation step of 5 min at 68 °C. The correct length of the amplicon was assessed via gel electrophoresis and the positive clones were inoculated in 5 mL LB medium containing 100 µg/mL ampicillin to grow an overnight culture, shaking at 37 °C. 4 mL of the culture were harvested by centrifugation and a plasmid Miniprep was performed using the PureYield Plasmid Miniprep System (Promega) according to manufacturer’s instructions. The pure plasmids were sent to sequencing at LGC genomics.

**Site directed mutagenesis**

The entire 8724 bp vector was amplified using back-to-back primers (primer sequences are shown in Supplemental Table 1) to insert the desired mutations. The first step was to remove the silent mutation at position c.882T>C; p.N294N using the primer pair FGFR3-SDMpos882T_F and FGFR3-pos876_R (primer sequences are shown in Supplemental Table S4). The resulting wild type vector was used as template for all subsequent PCRs to create the desired mutants. 10 ng of the template vector was added to the PCR as circular plasmid. Q5 High-Fidelity DNA Polymerase (NEB) was used at 0.3 U/µl in a 25 µl reaction in 1x Q5-reaction buffer supplemented with 1x GC-enhancer, 0.2 mM dNTPs (Biozym) and 0.4 µM of each primer. The PCR started with an initial heating step of 98 °C for 30 sec, followed by 25 cycles at 98 °C for 10 sec, an annealing step with variable temperatures (see Supplemental Table S4) for 15 sec, and 72 °C for 4.5 min, concluded by a final elongation step of 3 min at 72 °C in a T100 Thermal Cycler (BioRad). The correct length of the amplicon(s) was assessed via gel electrophoresis.

Next, PCR products forming partially nicked circular DNA were ligated for which 2 µl of the PCRs were used directly (without any purification step) in a 20 µl reaction supplemented with 1x T4 DNA Ligase Reaction Buffer (NEB) and 400 units T4 DNA Ligase (NEB). The reaction was mixed by pipetting and incubated over night at 16 °C. Afterwards, 10 µl of the ligation reaction were supplemented with 1x CutSmart buffer and 6 units of *DpnI* (NEB) and the methylated template plasmid was digested using an incubation time of 10 min at 37 °C followed by a heat inactivation of 20 min at 80 °C. Lastly, 10 µl of the *DpnI*-digest reaction were transformed into 50 µl of chemically competent *E. coli* XL1 Blue (Agilent) according to manufacturer’s instructions and plated on LB-agar containing 100 µg/mL ampicillin. Several clones were inoculated in 5 mL LB medium containing 100 µg/mL ampicillin to grow an overnight culture, shaking at 37 °C. 4 mL of the culture were harvested by centrifugation and a plasmid Miniprep was performed using the PureYield Plasmid Miniprep System (Promega) according to manufacturer’s instructions. The pure plasmids were sent to sequencing at LGC genomics. An overview of *FGFR3* mutant clones is listed in Supplemental Table S6.

**Cloning of positive control plasmid**

In order to delete the stop codon of *FGFR3* and to add an additional *AgeI* recognition sequence, the entire 8724 bp FGFR3(SP)-mGFP-5xGGS-FGFR3(EC-TM-IC)_pcDNA3.1/Hygro(+) vector (mGFP-FGFR3-WT) was amplified using the primer set XbaI-pcDNA3.1_F and FGFR3-w/o-stop-AgeI_R (primer sequences are shown in Supplemental Table S4). 10 ng of *XbaI* linearized vector was used as template for the PCR reaction using 0.3 U/µl Q5 High-Fidelity DNA Polymerase (NEB) in a 25 µl mix in 1x Q5-reaction buffer supplemented with 0.2 mM dNTPs (Biozym) and 0.4 µM of each primer. The PCR started with an initial heating step of 98 °C for 2 min, followed by 15 cycles at 98 °C for 10 sec, 66 °C for 15 sec, and 72 °C for 4 min 30 sec, followed by 10 cycles at 98 °C for 10 sec and 72 °C for 4 min 30 sec, concluded by a final elongation step of 3 min at 72 °C in a T100 Thermal Cycler (BioRad). The correct length of the amplicon was assessed via gel electrophoresis and the PCR amplicon was purified using the SeraMag Select (GE Healthcare) magnetic beads according to manufacturer’s instructions with a volume ratio of PCR to beads of 1:0.6. The vector-DNA was eluted in 37 µl nuclease-free water and the DNA concentration was determined using a Nanodrop 2000 instrument (Thermo Scientific).

mScarlet-I carrying a 5’ *AgeI* and a 3’ *XbaI* recognition sequence with additional 5’ bases to encode for the amino acids “GS” (to have a complete GGS-linker in between FGFR3 and mScarlet-I), was amplified using the primer set AgeI-GS-mScarlet_F and mScarlet-XbaI_R (primer sequences are shown in Supplemental Table S4) from our GRB2-mScarlet-I expression plasmid as template. 1 ng of circular plasmid was added to a PCR reaction using 0.3 U/µl Q5 High-Fidelity DNA Polymerase (NEB) in 25 µl total volume in 1x Q5-reaction buffer supplemented with 1x GC-Enhancer, 0.2 mM dNTPs (Biozym) and 0.4 µM of each primer. The PCR started with an initial heating step of 98 °C for 2 min, followed by 8 cycles at 98 °C for 10 sec, 60 °C for 15 sec, and 72 °C for 20 sec, followed by 20 cycles at 98 °C for 10 sec and 72 °C for 30 sec, concluded by a final elongation step of 3 min at 72 °C in a T100 Thermal Cycler (BioRad). The correct length of the amplicon was assessed via gel electrophoresis and the PCR amplicon was purified using the SeraMag Select (GE Healthcare) magnetic beads according to manufacturer’s instructions with a volume ratio of PCR to beads of 1:1. The DNA was eluted in 32 µl nuclease-free water and the DNA concentration was determined using a Nanodrop 2000 instrument (Thermo Scientific).

About 500 ng of PCR amplified vector DNA were digested with 20 units of the restriction enzymes *AgeI*-HF (NEB) and *XbaI* (NEB) in 1x CutSmart Buffer using an incubation time of 90 min at 37 °C followed by dephosphorylation using the Antarctic Phosphatase (NEB) in 1x Antarctic Phosphatase Buffer for 30 min at 37 °C, followed by a heat inactivation of 20 min at 65 °C. The digested linear vector was purified using the SeraMag Select (GE Healthcare) magnetic beads according to manufacturer’s instructions with a volume ratio of digest to beads of 1:0.8. The vector-DNA was eluted in 32 µl nuclease-free water and the DNA concentration was determined using a Nanodrop 2000 instrument (Thermo Scientific). Approximately 1 µg of bead-purified insert PCR amplicon was digested with 20 units of the restriction enzymes *AgeI*-HF (NEB) and *XbaI* (NEB) in 1x CutSmart Buffer using an incubation time of 90 min at 37 °C. After the digest, a buffer exchange was performed using the SeraMag Select (GE Healthcare) magnetic beads according to manufacturer’s instructions with a volume ratio of digest to beads of 1:1. The insert-DNA was eluted in 27 µl nuclease-free water and the DNA concentration was determined using a Nanodrop 2000 instrument (Thermo Scientific).

For ligation, 50 ng of digested vector-DNA were mixed with digested insert-DNA in a 1:3 vector-to-insert ratio in a 20 µl reaction supplemented with 1x T4 DNA Ligase Reaction Buffer (NEB) and 200 units T4 DNA Ligase (NEB). The reaction was mixed by pipetting and incubated first for 15 min at RT, then for 1 h at 16 °C and was lastly put on ice for 15 min. 9 µl of the ligation reaction were transformed into 50 µl of chemically competent *E. coli* NEB 10-beta (NEB) according to manufacturer’s instructions and plated on LB-agar containing 100 µg/mL ampicillin. Screening for positive clones was performed by colony PCR using the primer set FGFR3_pos2343_F and mScarlet-XbaI_R was used (primer sequences are shown in Supplemental Table S4). OneTaq® (NEB) polymerase was used at 0.025 U/µl in a 20 µl reaction in 1x standard buffer supplemented with 0.2 mM dNTPs (Biozym) and 0.5 µM of each primer. As template, the single colonies were dipped into the PCR master mix. The PCR started with an initial heating step of 94 °C for 5 min, followed by 30 cycles at 94 °C for 15 sec, 58 °C for 20 sec, and 68 °C for 50 sec, concluded by a final elongation step of 5 min at 68 °C. The correct length of the amplicon was assessed via gel electrophoresis and the positive clones were inoculated in 5 mL LB medium containing 100 µg/mL ampicillin to grow an overnight culture, shaking at 37 °C. 4 mL of the culture were harvested by centrifugation and a plasmid Miniprep was performed using the PureYield Plasmid Miniprep System (Promega) according to manufacturer’s instructions. The pure plasmids were sent to sequencing at LGC genomics.
